# Exploring the genomic diversity and antimicrobial susceptibility of *Bifidobacterium pseudocatenulatum* in the Vietnamese population to aid probiotic design

**DOI:** 10.1101/2021.04.22.441051

**Authors:** Hao Chung The, Chau Nguyen Ngoc Minh, Chau Tran Thi Hong, To Nguyen Thi Nguyen, Lindsay J. Pike, Caroline Zellmer, Trung Pham Duc, Tuan-Anh Tran, Tuyen Ha Thanh, Minh Pham Van, Guy E. Thwaites, Maia A. Rabaa, Lindsay J. Hall, Stephen Baker

**Affiliations:** Oxford University Clinical Research Unit, Ho Chi Minh City, Vietnam; The Wellcome Sanger Institute, Hinxton, Cambridge, United Kingdom; University of Cambridge School of Clinical Medicine, Cambridge Biomedical Campus, Cambridge, United Kingdom; Department of Medicine, University of Cambridge School of Clinical Medicine, Cambridge Biomedical Campus, Cambridge, United Kingdom; Centre for Tropical Medicine and Global Health, Nuffield Department of Clinical Medicine, Oxford University, Oxford, United Kingdom; Quadram Institute Biosciences, Norwich Research Park, Norwich, United Kingdom; Norwich Medical School, University of East Anglia, Norwich Research Park, Norwich, United Kingdom; Intestinal Microbiome, School of Life Sciences, ZIEL - Institute for Food & Health, Technical University of Munich, Freising, Germany

**Author notes:** Corresponding author: Dr. Hao Chung The, Department of Molecular Epidemiology, Oxford University Clinical Research Unit (OUCRU), 764 Vo Van Kiet, Ward 1, District 5, Ho Chi Minh City, Vietnam. equal contribution (Hao Chung The, Chau Nguyen Ngoc Minh).

**Keywords:** *Bifidobacterium* genomic, *Bifidobacterium pseudocatenulatum*, *Bifidobacterium* antimicrobial resistance, *Bifidobacterium* probiotic, genomic diversity, developing country, probiotic design, glycosyl hydrolase, exopolysaccharide

## Abstract

*Bifidobacterium pseudocatenulatum* is a member of the human gut microbiota, and has previously been used as a probiotic to improve gut integrity and reduce inflammatory responses. We showed previously that *B. pseudocatenulatum* was significantly depleted during dysenteric diarrhea, suggesting the organism may aid in recovery from diarrhea. Here, in order to investigate its probiotic potential, we aimed to assess the genomic diversity and predicted metabolic profiles of *B. pseudocatenulatum* found colonizing the gut of healthy Vietnamese adults and children. We found that the population of *B. pseudocatenulatum* from each individual was distinct and highly diverse, with intra-clonal variation attributed to gain or loss of carbohydrate utilizing enzymes. The *B. pseudocatenulatum* genomes were enriched with glycosyl hydrolases that target plant-based non-digestible carbohydrates (GH13, GH43), but not host-derived glycans. Notably, the exopolysaccharide biosynthesis region from organisms isolated from healthy children showed greater genetic diversity, and was subject to a high degree of genetic modification. Antimicrobial susceptibility testing revealed that the Vietnamese *B. pseudocatenulatum* were uniformly susceptible to beta-lactams, but exhibited variable resistance to azithromycin, tetracycline, ciprofloxacin and metronidazole. The genomic presence of *ermX* and *tet* variants conferred resistance against azithromycin and tetracycline, respectively; ciprofloxacin resistance was associated with mutation(s) in the quinolone resistance determining region (GyrA, S115 and/or D119). Our work provides the first detailed genomic and antimicrobial resistance characterization of *B. pseudocatenulatum* found in the Vietnamese population, which could inform the next phase of rational probiotic design.

**Importance:** *Bifidobacterium pseudocatenulatum* is a probiotic candidate with potential applications in several health conditions, but its efficacy is largely strain-dependent and associated with distinct genomic and biochemical features. However, most commercial probiotics have been developed by Western institutions, which may not have ideal efficacy when administered in developing countries. This study taps into the underexplored diversity of the organism in Vietnam, and provides more understanding to its lifestyles and antimicrobial susceptibility. These data are key for selecting an optimal probiotic candidate, from our established collection, for downstream investigations and validation. Thus, our work represents a model in identifying and characterizing bespoke probiotics from an indigenous population in a developing setting.

## Introduction

*Bifidobacterium* is a genus of Gram-positive non-spore forming anaerobic bacteria and among the most well-studied members of the human gut microbiota (1). These bacteria are among the major components of the gut microbiota and are transferred vertically from mothers to newborns. They are also measurably enriched in babies delivered vaginally, as compared to those delivered via caesarean section (2). Several health-promoting benefits are associated with *Bifidobacterium* colonization of the human gut. These benefits are associated with the production of secondary metabolites, immunomodulatory activities, and protection from infections (1, 3–5). The genus is composed of multiple human-adapted species, many of which colonize the gut during different life stages; this colonization pattern is largely dependent on the dominant carbohydrate sources available in the intestinal lumen (6). The saccharolytic lifestyle of *Bifidobacterium* can be observed by its ability to catabolize a wide variety of carbohydrates (from monosaccharides to complex plant-derived polysaccharides), which are ultimately channeled into a unique hexose metabolic pathway (“bifid shunt”) (7). This biochemical process permits the bacteria to generate more energy (in the form of ATP) from the same carbohydrate input, in comparison to the fermentative process found in other lactic acid bacteria (LAB) (7). Certain species and variants of *Bifidobacterium* are able to metabolize components of the early life diet, i.e. human milk oligosaccharides (HMO) present in breast milk, with *B. longum* subsp. *infantis* (8), *B. breve* (9), and *B. kashiwanohense* (10) enriched in the intestines of breast-fed infants. After weaning, *Bifidobacterium* ceases to predominate in the gut (11), and only species that can thrive on complex dietary carbohydrates are able to flourish. These species include *B. longum* subsp. *longum* (12, 13), *B. adolescentis*, and *B. pseudocatenulatum (Bp)*.

*Bp* is less well characterized than other *Bifidobacterium* species, but is associated with several health benefits. Expansion of *Bp* in the gut microbiome was associated with successful weight loss in obese children in China following ∼100-day fiber-rich dietary (FRD) interventions (14, 15). According to a recent clinical trial, *Bp* was also among the enriched short-chain fatty-acid (SCFA)-producing gut commensals in type-2 diabetes patients receiving FRD interventions (16). Additionally, experimental evidence has demonstrated that supplementation with a *Bp* probiotic (CECT 7765) in obese mice led to improved metabolic responses (lowering serum cholesterol, triglyceride, and glucose concentrations) (17) and reduced pro-inflammatory cytokines (IL-17A and TNF-α) (18). A further trial in obese Spanish children with insulin resistance demonstrated that treatment with the same probiotic resulted in a comparable improvement in inflammatory status (19). Moreover, recent studies demonstrated that oral administration of *Bp* enhanced gut barrier integrity and alleviated bacterial translocation in mice with induced liver damage (20, 21).

In a recent microbiome study, we found that *Bp* was consistently depleted in the gut microbiomes of Vietnamese children suffering from dysenteric (mucoid and/or bloody) diarrhea(22). This association remained significant regardless of etiological agent. Dysenteric diarrhea is associated with heightened inflammation, and we hypothesized that *Bp* may be beneficial in reducing inflammation-associated conditions and accelerating recovery of the gut microbiota following diarrhea. The efficacy of *Bp* as a probiotic is largely strain-dependent and associated with distinct genome composition and biochemical profile (23). However, most commercially available probiotics have been developed by Western institutions, which may not have ideal gut colonization and efficacy when administered in tropical or developing countries. Therefore, aiming to generate data to support the development of a candidate *Bp* probiotic suitable for use in treating/preventing dysenteric diarrhea(24), we assessed the genetic diversity of *Bp* colonizing the guts of healthy Vietnamese children and adults. Here, we defined the genetic diversity, predicted biochemical profile, and antimicrobial susceptibility of the *Bp* population in the Vietnamese population. These data are key for selecting an optimal probiotic candidate for downstream investigations and validation, with particular consideration for its potential uses in LMICs in Southeast Asia.

## Results

### The abundance of Bifidobacterium pseudocatenulatum in the Vietnamese population

In order to investigate the distribution and diversity of *Bifidobacterium* spp. in the gut microbiota of Vietnamese adults and children, we extracted total DNA from fecal samples collected from 42 healthy Vietnamese individuals (21 children and 21 adults) and subjected them to shotgun metagenomic sequencing. All recruited children were aged between 9 and 59 months (median: 23 [interquartile range: 9 – 37] months) and had been weaned onto a solid food diet for at least three months. All recruited adults were aged between 25 and 59 years (median: 35 years) and reported having an omnivorous diet.

Taxonomic profiling from the microbiome data demonstrated that *Bifidobacterium* species were more abundant in children compared to adults (median relative abundances; 8.0% [3.8 – 19.7], and 1.2% [0.3 – 3.4], respectively) (Fig 1). Specifically, we found that *Bp* was the most prevalent *Bifidobacterium* species in the adults (mean = 1.1%) and the second-most prevalent *Bifidobacterium* species in children (mean = 2.9%, after *B. longum*).

**Figure 1.**
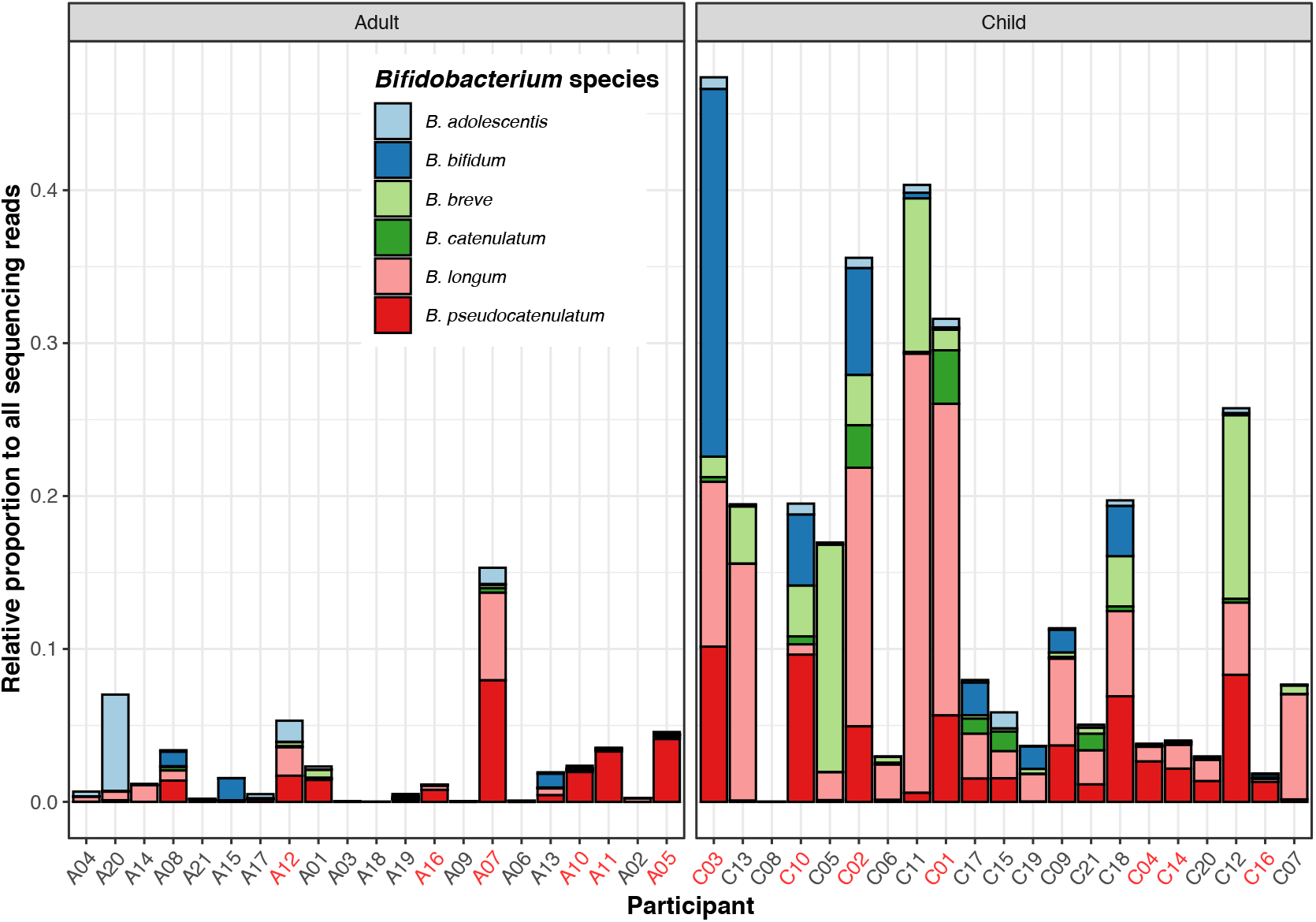
Abundance of *Bifidobacterium* species in the gut microbiomes of a Vietnamese population. The figure displays the relative abundance of *Bifidobacterium* (calculated as percentage of reads classified as *Bifidobacterium* by Kraken, relative to each sample’s total sequencing reads) in the gut microbiomes of adult (left) and child (right) participants (n=21 for each group). The sample names in each group are ordered based on the participants’ age, in increasing order. Samples labelled in red denote successful culture of laboratory-confirmed *B. pseudocatenulatum*. Different *Bifidobacterium* species are colored as in the key.

### The phylogenetic distribution of Vietnamese Bifidobacterium pseudocatenulatum

We selected fecal samples from participants with a relative *Bp* abundance >0.1% (n=16) to isolate *Bifidobacterium*. In total, we isolated 185 individual organisms with a colony morphology indicative of *Bifidobacterium*. Among these, 49 isolates (from 7 children and 6 adults) were *Bp* according to MALDI-TOF bacterial identification and full-length sequencing of the 16S rRNA (Fig 1). These 49 individual organisms were subjected to whole genome sequencing (WGS). A preliminary phylogenetic reconstruction using a core-genome alignment segregated the organisms into two distinct lineages (Fig S1). The majority of isolates (n=45) clustered with two *Bp* reference genomes (DSM20438 and CECT_7756) within the major lineage. The remaining four isolates (C01_H5, C01_D5, C01_C5, C02_A8) formed a separate cluster that was distantly related to the major lineage. Further interrogation and comparison with *B. catenulatum* and *B. kashiwanohense* genomes confirmed that these four isolates belonged to the *B. catenulatum* spp. complex (Fig S2).

Refined phylogenetic reconstruction of the 45 *Bp* genomes identified 13 phylogenetic clusters (PCs) and three singletons (C14_S, A05_S, A16_S) (Fig 2), all of which were supported by high bootstrap values (>80%). For ease of nomenclature, these PCs and singletons were collectively called PCs. Each PC was defined by close genetic relatedness (negligible branch lengths), and each contained organisms isolated exclusively from a single individual. However, isolates recovered from each sampled participant were either solely restricted to one PC (6/11 participants) or distributed across two PCs (5/11). Moreover, when multiple PCs were detected within the same individual, they were generally not monophyletic (except for C16). These data suggest that the *Bp* population within each individual is highly diverse, and PCs could not be distinguished based on the age of the participant (child versus adult).

**Figure 2.**
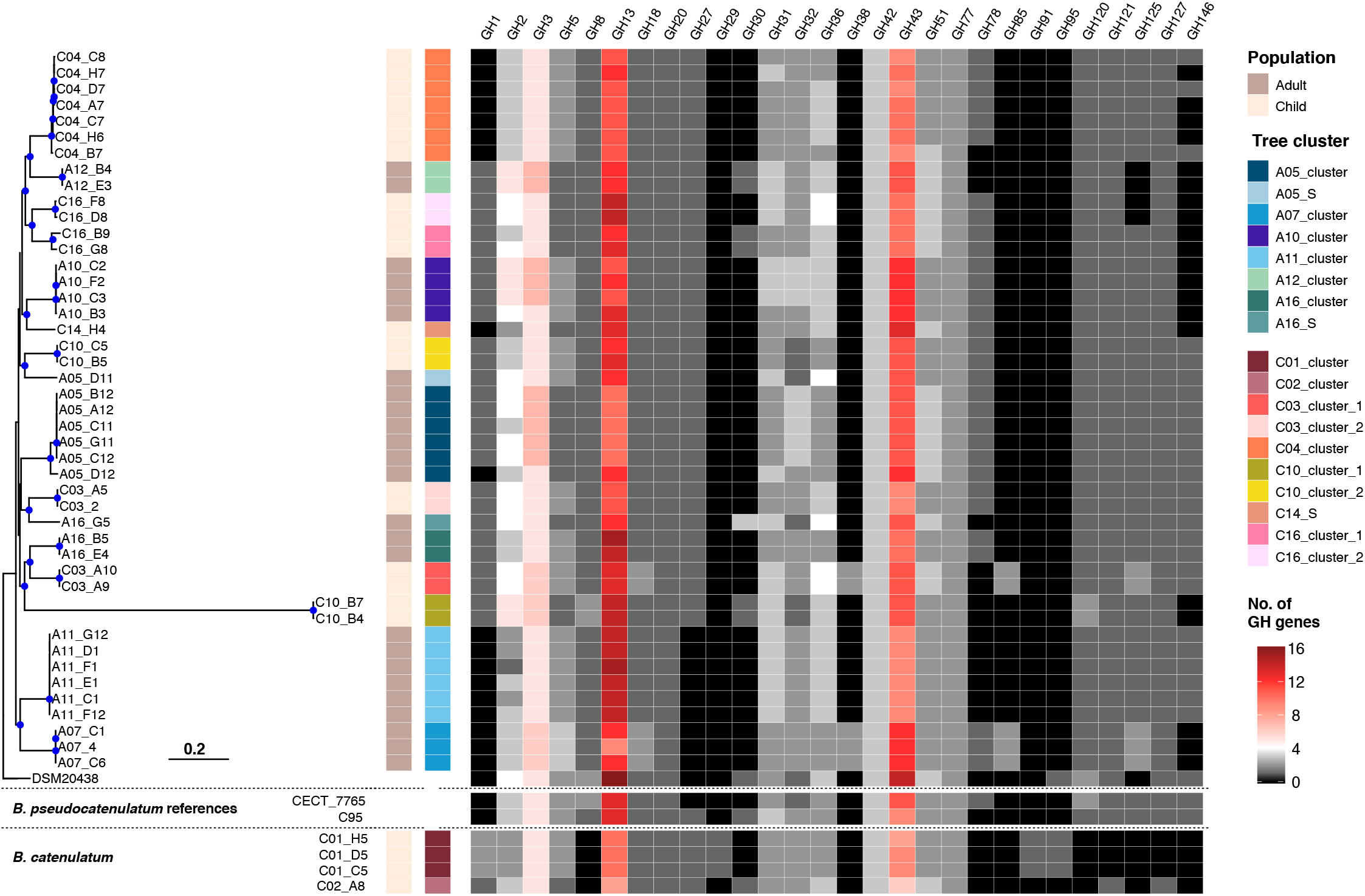
The phylogeny and glycosyl hydrolase profile of Vietnamese *Bifidobacterium pseudocatenulatum*. The maximum likelihood phylogeny of 45 *B. pseudocatenulatum* isolated in this study was constructed from a recombination-free alignment following read mapping (See Methods). The tree is rooted using the reference DSM20438 as an outgroup, and blue filled circles denote bootstrap values greater than 70 at the internal nodes. Other *B. pseudocatenulatum* references (CECT_7765 and the Chinese isolate C95) and *B. catenulatum* isolated in this study are included for comparison. The columns to the right of the phylogeny show the metadata associated with each taxon, including population, tree cluster nomenclature, and the numbers of gene belonging to each defined glycosyl hydrolase family (GH1, GH2, GH3, GH5, GH8, GH13, GH18, GH20, GH27, GH29, GH30, GH31, GH32, GH36, GH38, GH42, GH43, GH51, GH77, GH78, GH85, GH91, GH95, GH120, GH121, GH125, GH127, GH146). The quantity of these genes is denoted according to the key. The horizontal scale bar denotes the number of substitutions per site.

### In silico prediction of carbohydrate utilization

*Bifidobacterium* spp. are renowned for their ability to utilize a diverse range of carbohydrates, which contribute to the functional integrity of the human gut. We focused on identifying the repertoire of carbohydrate utilizing enzymes (CAZymes) within the *Bp* genome sequences to predict the carbohydrate metabolic capacity of each isolate. Among 4,333 gene families in the pangenome, 233 were determined to be CAZymes. These included 126 glycosyl hydrolases (GH), 97 glycosyltransferases (GT), two carbohydrate esterases, and two carbohydrate binding motif (CBM) containing proteins. As GHs catalyze the breakdown of glycosidic bonds, they are essential for the assimilation of complex glycans. We mapped the presence of all GH genes in each isolate of the *Bp* (45 Vietnamese and 7 reference isolates) and *B. catenulatum* (C01 and C02 clusters) collections. Genes pertaining to GH23 and GH25 were excluded from interpretation as they participate specifically in the recycling of the peptidoglycan in the bacterial cell wall.

Thirty-four GH genes were classified as core (present in all 52 *Bp* genomes), while accessory GH genes were more enriched in ≤10 genomes (Fig S3). The predominant GH families identified were GH13 (median of 12 copies per isolate) and GH43 (median of 10.5 copies per isolate), followed by GH3

(median of 5 copies per isolate) (Fig 2). GH13 mainly catalyze the hydrolysis of α-glucosidic linkages (in resistant starch and α-glycans), while GH3 is involved in the assimilation of cellobiose and cellodextrin. GH43 includes a diverse range of α-L-arabinofuranosidase, β-xylosidase, and xylanase involved in the degradation of hemicellulose, arabinogalactan, arabinan, and arabinoxylan. In contrast, GH targeting host-derived glycans (GH29, GH33, GH35, and GH95) were not detected this *Bp* collection. These data, coupled with the variable presence of other GH families (GH31, GH32, GH36, GH42, GH51, and GH77), signify a tropism for dietary starch and fiber in the catabolism of these organisms.

### Genomic variation within the phylogenetic cluster

Given that the phylogeny was generated from the core genome, we considered that isolates belonging to the same PC have limited variation in the core genome but may have substantial variation within their accessory genomes. This genetic variation may underlie major phenotypic differences within a single clone. We investigated the presence/absence of accessory genes in the pangenome of each PC and found that the distribution of such variation was not uniform. Inter-strain variation was found to be minimal for 7/13 non-singleton PCs (≤ 26 differences) (Fig 3A). This limited genetic diversity may be attributed to insufficient sampling coverage; however, more intensive sampling, as demonstrated in the A11_cluster (six isolates), still resulted in low variation in the pangenome. These differences in intra-clonal gene content were typically associated with genes encoding carbohydrate transport (ABC transporter and permease) and utilization proteins (GH, GT, esterase), or were of unknown function (6 – 25 hypothetical proteins per PC). Alternatively, the bimodal distribution observed in the A05_cluster, A07_cluster, and A10_cluster demonstrates that while most organisms share limited variation in gene content (∼12 genes), outlier organisms may carry a distinct accessory genome. This observation resulted in sizeable differences when comparing the outlier to the remainders in each PC. For example, the accessory genome of A05_D12 (A05_cluster) differed from that of the remaining five isolates by >100 genes. This composition of genes arose from a recombination event (spanning 28 kbp from 2,126,733 to 2,154,962 in the DSM20438 chromosome and containing multiple ABC transporters and GHs) and the gain of an IS3-mediated region (ABC transporters and β-glucosidase), which distinguished A05_D12 from the other isolates.

**Figure 3.**
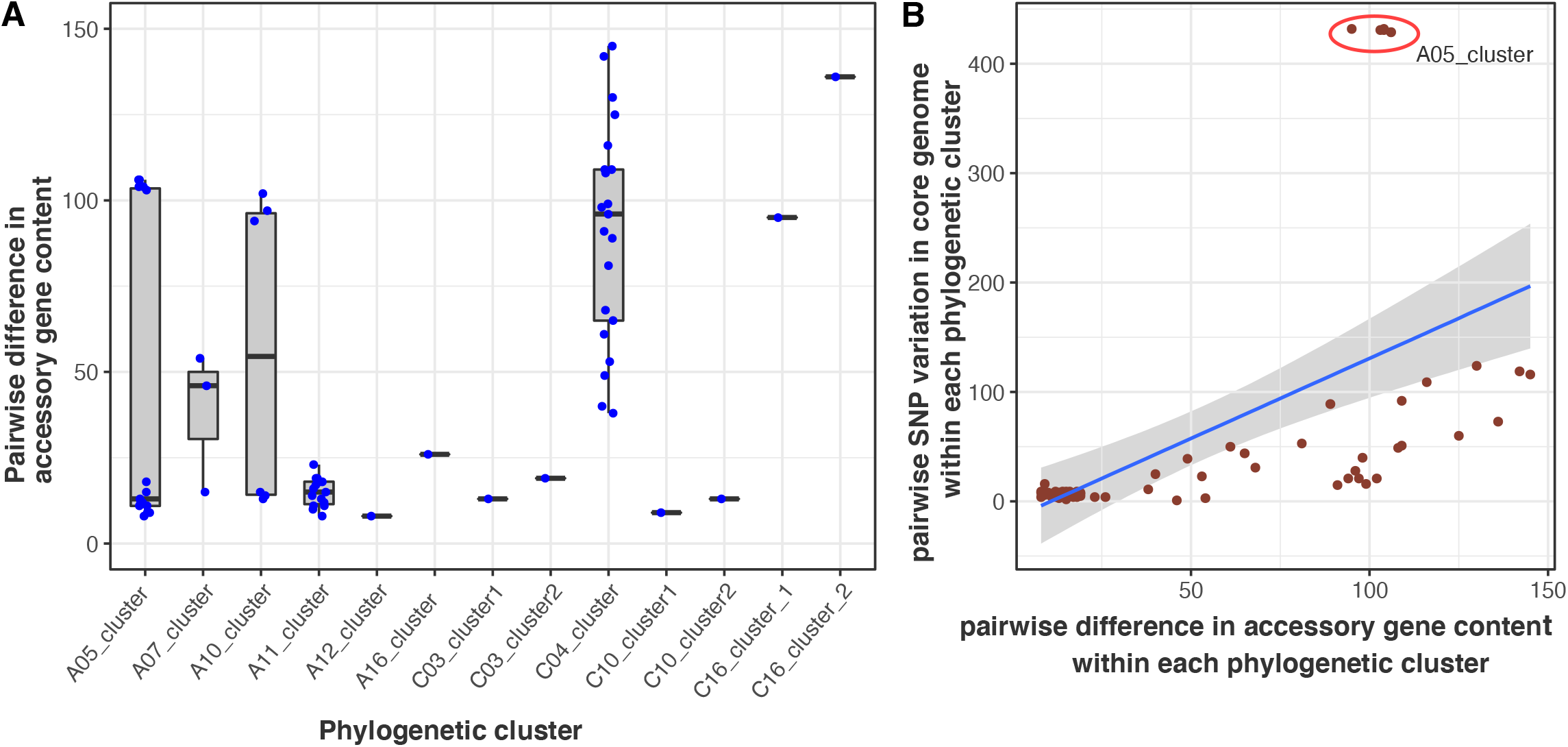
Variation in the accessory genomes of *Bifidobacterium pseudocatenulatum*. (A) The panel depicts the distribution of pairwise differences in the accessory gene content (counted as presence/absence, represented as blue circles) of isolates within each defined tree cluster. For each boxplot, the upper whisker extends from the 75^th^ percentile to the highest value within the 1.5 * interquartile range (IQR) of the hinge, and the lower whisker extends from the 25^th^ percentile to the lowest value within the 1.5 * IQR of the hinge. (B) The panel displays the positive correlation between intra-clonal pairwise differences in the accessory gene content (x-axis) and intra-clonal pairwise variation of single nucleotide polymorphisms (SNPs) in the core genome. The red circle indicates the outlier, A05_cluster_1.

Most noticeably, the C04_cluster contained a wide distribution of pairwise-differences across the pangenome, indicating that each isolate possessed a moderately divergent accessory genome. To visualize the magnitude of lateral gene transfer, we separately reconstructed the phylogeny of the C04_cluster (Fig S4). Branch lengths indicative of significant divergence, coupled with a high frequency of gene acquisition and gene loss events, demonstrate that the C04 *Bp* population has undergone extensive clonal expansion. This micro-evolution underlies a diversifying metabolic potential, as exemplified by the concurrent acquisition of a GH78 (rhamnosidase) and deletion of a GH51 (arabinosidase) in a specific monophyletic cluster (Fig 2 and S4). Notably, a novel ribulose-5P-3-epimerase gene was acquired in the most recent common ancestor (MRCA) of C04_A7 and C04_D7. This enzyme bilaterally converts ribulose-5-P to xylulose-5P, a key intermediate in the *Bifidobacterium* specific hexose metabolic pathway (bifid shunt) (7), thus potentially facilitating a greater energy harvest. We aimed to identify the genetic origin of the acquired elements (by BLAST to public database) and found that *B. kashiwanohense* and *B. adolescentis* were the most likely sources.

We hypothesized that the extent of intra-PC variation in the accessory genome was dependent on the evolutionary timeframe of the PC, which is reflected in the number of core genome SNPs the PC has accumulated since its MRCA. Within the examined PCs, the median of pairwise recombination-free SNPs was 9 (IQR [13 – 49]), while the median variation in the pangenome was 24 genes (IQR [13 – 97]). As shown in Figure 3B, the pairwise difference in the accessory genome partially correlated with the pairwise SNP distance (Pearson’s r = 0.54). An outlier to this trend was in the A05_cluster (retrieved from a 59-year-old female), in which A05_D12 was >400 SNPs away from the remaining five closely related isolates, albeit with only ∼100 gene differences in the accessory genome. These findings suggest that as the *Bp* population undergoes a prolonged period of within-host evolution and expansion, its pan-genome may expand through increasing horizontal gene transfer (HGT).

### Genomic differences in Bifidobacterium pseudocatenulatum originating from children and adults

The differing physiologies and diets of children and adults create distinct niches in which *Bifidobacterium* can adapt, and such adaptation may be reflected by genomic variation. An exploratory analysis of 21 representative isolates (10 from adults, 11 from children) identified 42 genes with differing abundance between *Bp* originating from children and adults. Of these 42 genes, 13 were of unknown function. Eight genes (4 glycosyltransferases, a polysaccharide export *rfbX*, an O-acetyltransferase, a reductase, and *fhiA*) were more frequently present in bacteria retrieved from adults (6/10 representative clusters), compared to those from children (1/11 representative clusters) (Fisher’s Exact test, *p*=0.024). These genes formed the core component of the exopolysaccharide (EPS) biosynthesis cluster of the reference strain DSM20438 (Fig 4). We characterized this genomic region of all representative *Bp* in detail and confirmed that the EPS region in organisms from adults was more similar to that of DSM20438 (Table 1). In organisms isolated from both adults and children, the EPS cluster was subject to frequent modification, with integrations of genes derived from the EPS region of other *Bifidobacterium* species, such as *B. longum* and *B. kashiwanohense*. Notably, the rhamnose precursor biosynthesis genes (*rmlABC*) were present in isolates of three clusters (C03_cluster_1, A07_cluster, A16_cluster), which predicts the incorporation of rhamnose or rhamnose-derived sugar in the EPS structure (25). Organisms of distantly related PCs occasionally shared comparable EPS regions, as observed in A05_S and the A11_cluster. Specifically, the EPS region of C03_cluster_1 was similar to that of *Bp* CECT_7765, which has been developed as a probiotic candidate to alleviate inflammatory responses in patients with cirrhosis and obesity in Spain(19, 26).

**Table 1.**
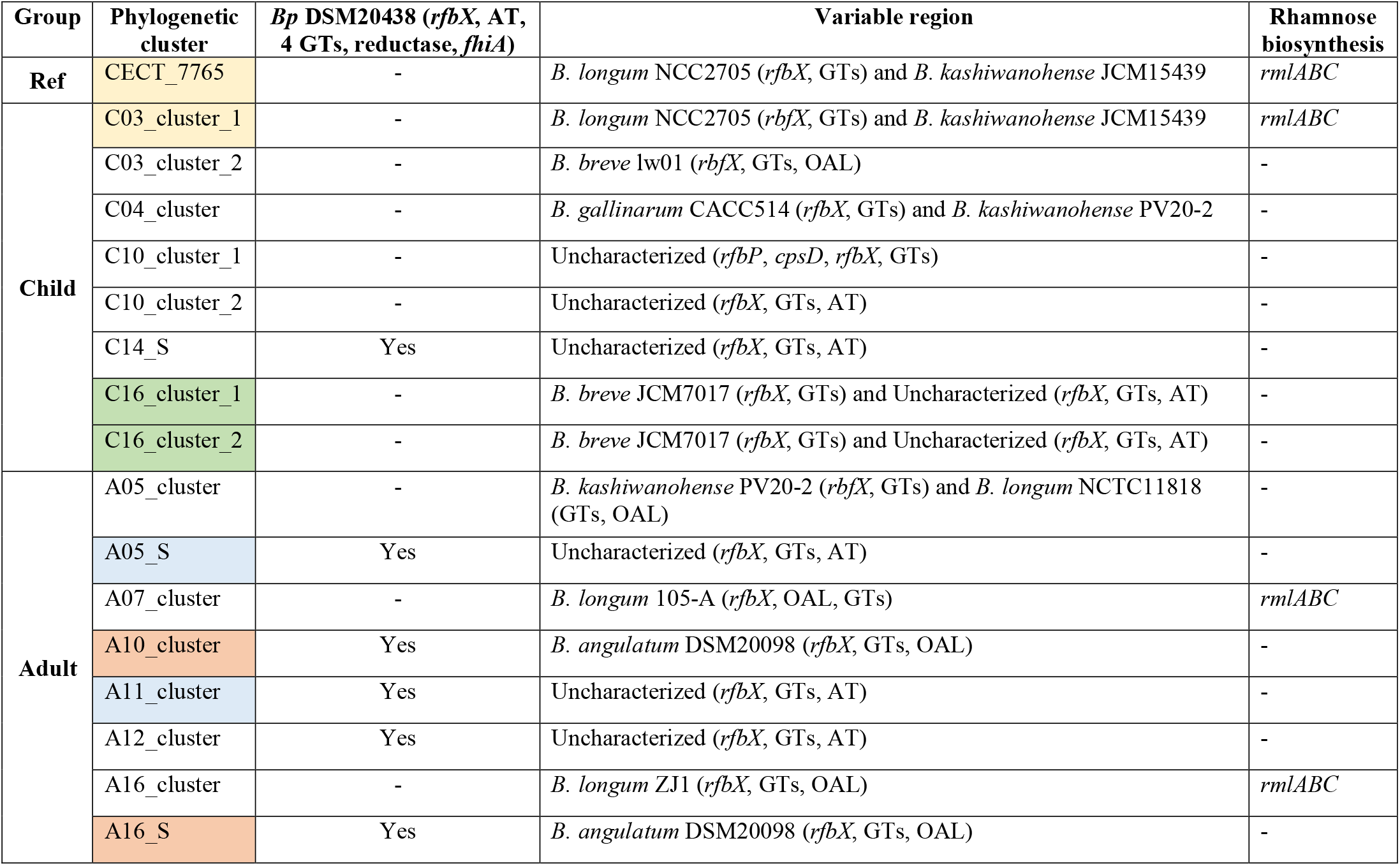
Exopolysaccharide biosynthesis of *B. pseudocatenulatum*. GT: glycosyltransferase; AT: acyl-transferase; OAL: O-antigen ligase; *rfbX*: O-antigen transporter. *rfbP, cpsD*: Priming glycosyltransferase. Bp: *B. pseudocatenulatum*. Isolates with the same shading colour share similar EPS biosynthesis cluster.

| Group | Phylogenetic cluster | *Bp* DSM20438 (*rfbX*, AT, 4 GTs, reductase, *fhiA*) | Variable region | Rhamnose biosynthesis |
| --- | --- | --- | --- | --- |
| Ref | CECT_7765 | - | *B. longum* NCC2705 (*rfbX*, GTs) and *B. kashiwanohense* JCM15439 | *rmlABC* |
| Child | C03_cluster_1 | - | *B. longum* NCC2705 (*rfbX*, GTs) and *B. kashiwanohense* JCM15439 | *rmlABC* |
|  | C03_cluster_2 | - | *B. breve* lw01 (*rfbX*, GTs, OAL) | - |
|  | C04_cluster | - | *B. gallinarum* CACC514 (*rfbX*, GTs) and *B. kashiwanohense* PV20-2 | - |
|  | C10_cluster_1 | - | Uncharacterized (*rfbP*, *cpsD*, *rfbX*, GTs) | - |
|  | C10_cluster_2 | - | Uncharacterized (*rfbX*, GTs, AT) | - |
|  | C14_S | Yes | Uncharacterized (*rfbX*, GTs, AT) | - |
|  | C16_cluster_1 | - | *B. breve* JCM7017 (*rfbX*, GTs) and Uncharacterized (*rfbX*, GTs, AT) | - |
|  | C16_cluster_2 | - | *B. breve* JCM7017 (*rfbX*, GTs) and Uncharacterized (*rfbX*, GTs, AT) | - |
| Adult | A05_cluster | - | *B. kashiwanohense* PV20-2 (*rfbX*, GTs) and *B. longum* NCTC11818 (GTs, OAL) | - |
|  | A05_S | Yes | Uncharacterized (*rfbX*, GTs, AT) | - |
|  | A07_cluster | - | *B. longum* 105-A (*rfbX*, OAL, GTs) | *rmlABC* |
|  | A10_cluster | Yes | *B. angulatum* DSM20098 (*rfbX*, GTs, OAL) | - |
|  | A11_cluster | Yes | Uncharacterized (*rfbX*, GTs, AT) | - |
|  | A12_cluster | Yes | Uncharacterized (*rfbX*, GTs, AT) | - |
|  | A16_cluster | - | *B. longum* ZJ1 (*rfbX*, GTs, OAL) | *rmlABC* |
|  | A16_S | Yes | *B. angulatum* DSM20098 (*rfbX*, GTs, OAL) | - |

**Figure 4.**
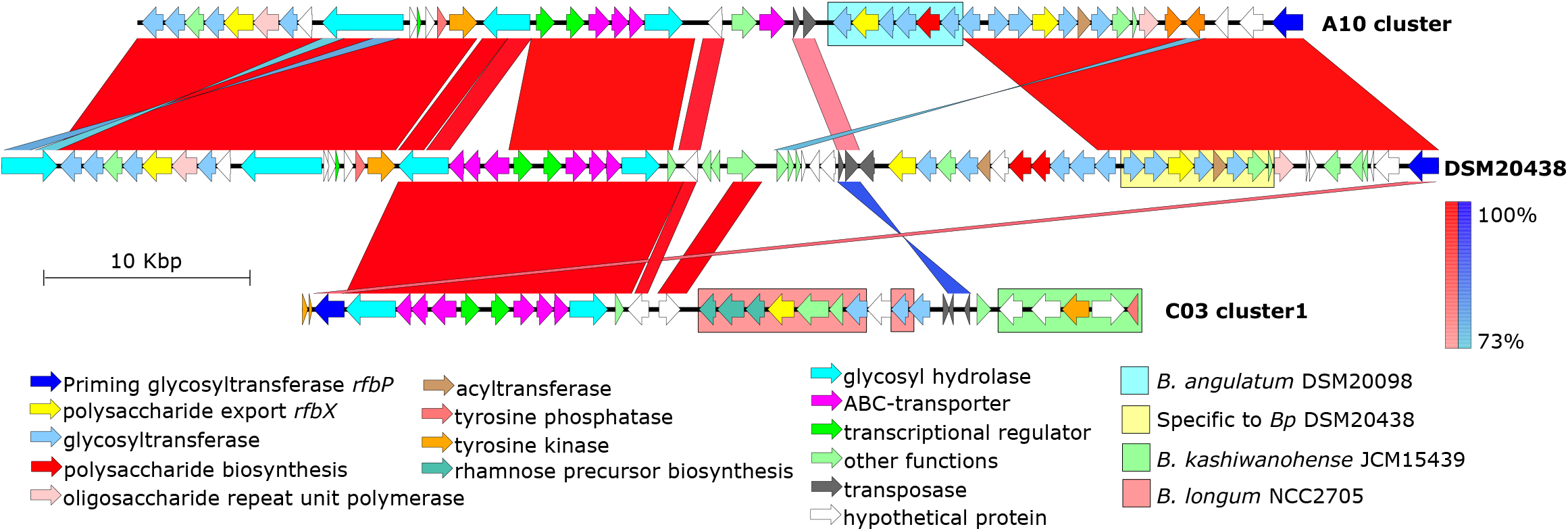
Comparison of the exopolysaccharide (EPS) biosynthesis genomic region from exemplar *Bifidobacterium pseudocatenulatum* genomes. Each arrow represents a predicted gene, with its function colored as in the keys. The blocks connecting two genomes indicate regions with high nucleotide similarity, either as synteny (red) or inversion (blue), with the color intensity corresponding to the degree of nucleotide similarity. The colored boxes denote regions homologous to that found in specific *Bifidobacterium* species.

Two additional genes were found to be enriched in *Bp* isolated from adults (8/10), compared to that from children (3/11) (Fisher’s Exact test, *p*=0.03). These two tandem genes (BBPC_RS09395 and BBPC_RS08115 in the reference DSM20438) both encode for GH43. RS09395 is a large multi-domain protein (>2,000 aa) and consists of three GH43 subunits. Among these, two subunits shared >70% nucleotide identity with the α-L-arabinofuranosidases, *arafB* (BLLJ_1853, GH43_22) and *arafE* (BLLJ_1850, GH43_34), of *B. longum* JCM1217, which encode for degradative enzymes targeting the arabinan backbone and arabinoxylan, respectively (27). The remaining GH43_22 subunit of RS09395 showed no ortholog in *B. longum* and shared 65% amino acid identity to that in *B. catulorum*. In contrast, RS08115 was smaller (∼1,000 aa) and shared 65% nucleotide identity with *arafA* (BLLJ_1854, GH43_22) of *B. longum* JCM1217, known to specifically degrade arabinogalactan (28). Bioinformatic analyses predicted that both RS09395 and RS08115 were secreted and bound to the bacterial cell membrane, due to the presence of N-terminal signal peptide and transmembrane motifs. These data suggest that these two enzymes synergistically degrade arabinoglycan, releasing degradants (i.e. L-arabinose) into the extracellular milieu and contributing to cross-feeding with other members of the gut microbiota.

### Antimicrobial susceptibility of representative Bifidobacterium pseudocatenulatum

To better evaluate their suitability for probiotic design, specifically to assess if they can be formulated along with antimicrobial treatments, we subjected the isolated *Bp* to antimicrobial susceptibility profiling. We reported a broad range of MICs against ceftriaxone, amoxicillin/clavulanate, ciprofloxacin, azithromycin, tetracycline, and metronidazole for 19 representative isolates (17 *Bp*, 2 *B. catenulatum*) (Table 2). Notably, the MICs for ceftriaxone (≤1.5 µg/mL) and amoxicillin/clavulanate (≤0.25µg/mL) were consistently low, likely indicating that all tested *Bifidobacterium* would be susceptible to these β-lactams during antimicrobial therapy. In contrast, susceptibility against the remaining antimicrobials was more variable, as evidenced by their broader MIC ranges. The highest MICs for ciprofloxacin (32 µg/mL) and metronidazole (256µg/mL) were observed in 57% (11/19) of tested *Bifidobacterium* isolates (Table 2). Concurrent non-susceptibility against ciprofloxacin (MIC = 32 µg/mL), azithromycin (MIC = 256 µg/mL), and metronidazole (MIC = 256 µg/mL) was observed in four isolates.

**Table 2.** Summary of E-test results for 21 *Bifidobacterium* isolates (19 representative *B. pseudocatenulatum* and 2 controls). CRO: Ceftriaxone (30µg); CIP: Ciprofloxacin (5µg); AMC: amoxicillin/clavulanic acid (30µg); AZM: azithromycin (15µg); TET: tetracycline (30µg); MTZ: metronidazole (5µg).

| Anti-microbial | Number of strains with MIC (µg/mL) |  |  |  |  |  |  |  |  |  |  |  |  |  |  |  |  |  |  |  |  |  |  |
| --- | --- | --- | --- | --- | --- | --- | --- | --- | --- | --- | --- | --- | --- | --- | --- | --- | --- | --- | --- | --- | --- | --- | --- |
|  | ≤0.047 | 0.064 | 0.094 | 0.125 | 0.19 | 0.25 | 0.38 | 0.5 | 0.75 | 1 | 1.5 | 2 | 3 | 4 | 8 | 12 | 16 | 24 | 32 | 48 | 64 | 128 | >256 |
| CRO |  |  | 2 | 4 | 2 | 4 | 1 | 3 | 2 |  | 1 |  |  |  |  |  |  |  |  |  |  |  |  |
| CIP |  |  |  |  |  |  |  | 1 | 1 | 3 | 1 | 2 |  |  |  |  |  |  | 11 |  |  |  |  |
| AMC | 2 | 2 | 4 | 7 | 2 | 2 |  |  |  |  |  |  |  |  |  |  |  |  |  |  |  |  |  |
| AZM | 1 |  | 1 |  | 1 | 1 | 1 | 2 | 2 | 1 |  |  | 1 |  | 1 |  |  |  |  | 1 |  | 1 | 5 |
| TET |  |  |  |  |  |  | 2 | 1 | 4 | 4 |  | 2 | 1 | 1 |  |  |  | 1 |  | 2 | 1 |  |  |
| MTZ |  |  |  |  |  |  |  |  |  |  |  | 1 |  |  |  | 1 | 1 | 1 |  | 2 | 1 | 1 | 11 |

We examined the correlation between MIC values and inhibitory zone diameters (IZD) for the six aforementioned antimicrobials (Figure 5). The narrow range of recorded values for amoxicillin/clavulanate (0.047 - 0.25 µg/mL, 36 – 48 mm) and ceftriaxone (0.094 - 1.5 µg/mL, 28 – 40 mm) resulted in a weak to modest negative correlation (Kendall’s r ≥ −0.5). Such correlation appeared to be stronger for azithromycin, tetracycline, and ciprofloxacin (Kendall’s r ≤ −0.7), likely owing to a wider range of MIC and IZD values. Specifically, for ciprofloxacin, an MIC value of 32 µg/mL corresponded with an IZD of 6 mm (no killing zone), while the remaining MIC (0.5 – 2 µg/mL) corresponded with IZDs >18 mm. In contrast, a poorer correlation was observed with metronidazole despite presenting a similarly broad range of MIC values, such that an IZD of 6 mm corresponded with a wide range of MICs (12 – 256 µg/mL). These results showed that, with exception of metronidazole, both E-test and disc diffusion methods produce robust and consistent interpretations for antimicrobial susceptibility in these *Bifidobacterium*.

**Figure 5.**
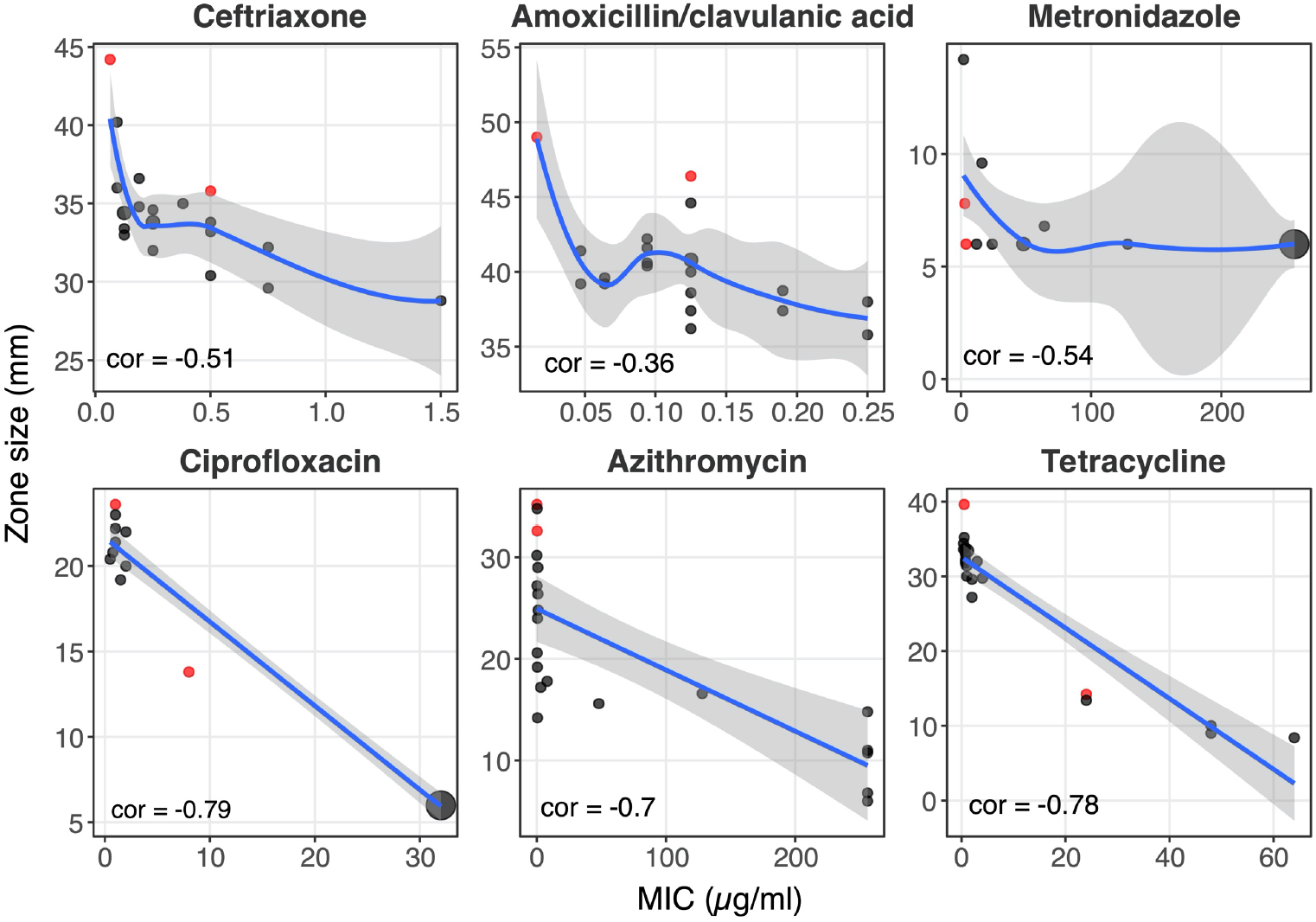
Correlation between E-test and disc diffusion methods in antimicrobial susceptibility testing of *Bifidobacterium*. Each panel represents a tested antimicrobial, with the x-axis and y-axis denoting the results of E-test (minimum inhibitory concentration in µg/ml) and disc diffusion (inhibitory zone diameter in mm) approaches. Controls (*B. pseudocatenulatum* DSM20438 and *B. longum* NCIMB 8809) are colored in red, while tested *Bifidobacterium* isolated in this study are colored in grey. The circle size is proportional to the number of isolates bearing the same MIC and IZD, and the largest circles in metronidazole and ciprofloxacin panels correspond to eleven isolates. All correlation scores are calculated using Kendall’s correlation. LOESS regression is shown for ceftriaxone, amoxicillin/clavulanic acid, and metronidazole (cor > −0.7), while linear regression is shown for ciprofloxacin, azithromycin, and tetracycline (cor ≤ −0.7).

### Antimicrobial resistance genetic determinants in Bifidobacterium pseudocatenulatum

We lastly sought to investigate potential mechanisms of antimicrobial resistance (AMR) in the sequenced *Bifidobacterium*. As resistance to metronidazole is complex and typically attributed to altered metabolism (29), we only focused on the genetic determinants for resistance against tetracycline, azithromycin, and ciprofloxacin. Screening against a curated database of acquired AMR genes revealed the presence of *tetO* and *ermX* in our isolates. The presence of *tetO* correlated significantly with elevated MIC and decreased IZD against tetracycline, while the presence of *ermX* was associated with a decrease in azithromycin IZD (Figure 6). As ciprofloxacin resistance is commonly induced by mutations in the quinolone resistance determining region (QRDR) on bacterial topoisomerases (30–32), we screened *gyrA, gyrB, parC*, and *parE* in the assembled *Bifidobacterium* genomes to identify nonsynonymous mutations in the QRDR. This analysis detected the presence of non-synonymous mutations in *gyrA*. These included single mutations such as S115F (n=5), S115V (n=1), S115Y (n=1), and D119G (n=3), as well as a double mutation, S115F - D119G (n=1). Upon classifying the isolates based on the presence of the aforementioned mutations, we found that their presence significantly correlated with elevated MIC (32 µg/mL) and reduced IZD (6 mm) (Figure 6). These data indicate that the presence of *tetO, ermX*, and specific mutations in *gyrA* in our *Bifidobacterium* collection explain non-susceptibility against tetracycline, azithromycin, and ciprofloxacin, respectively.

**Figure 6.**
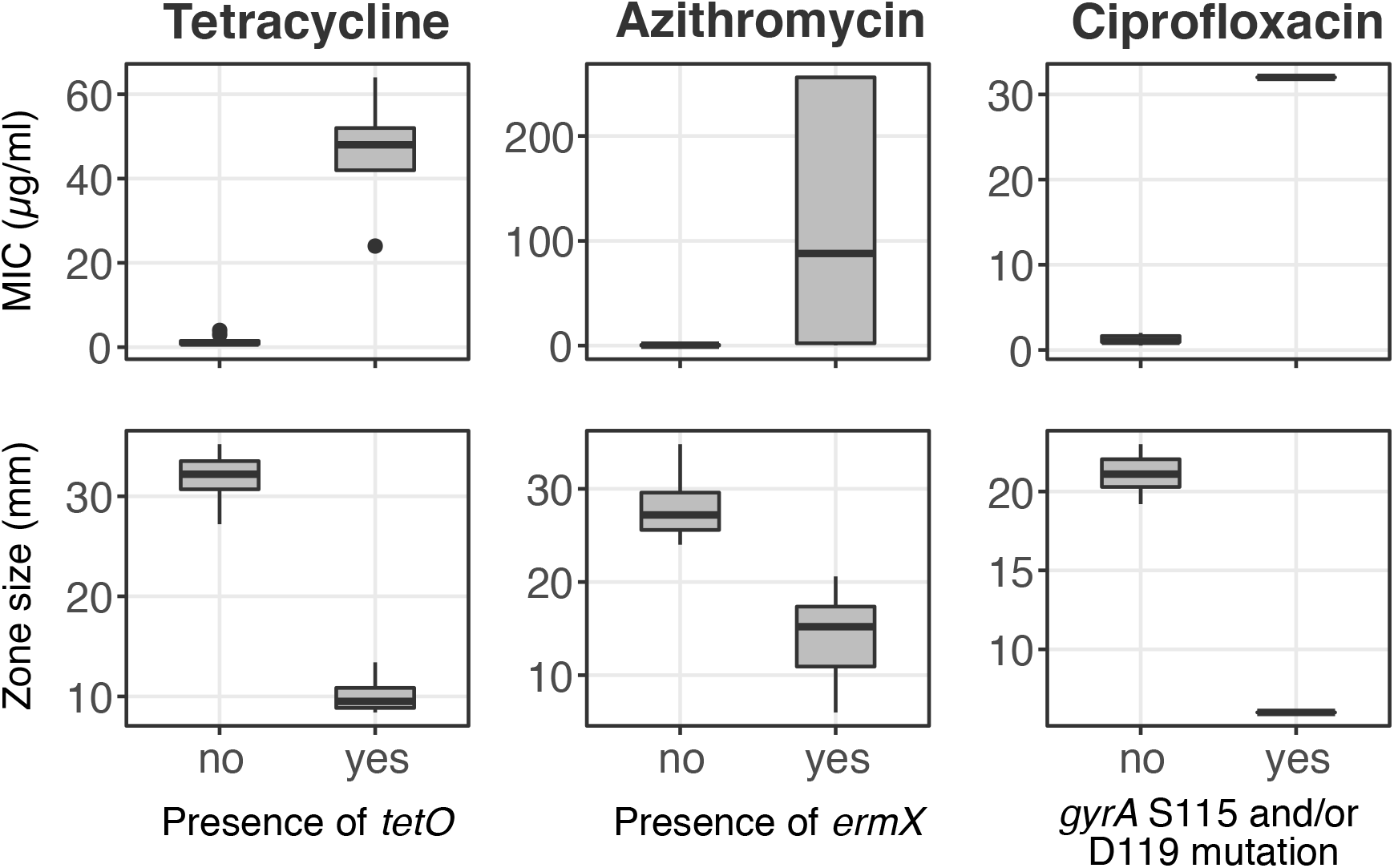
Association between resistance determinants and antimicrobial testing results in *Bifidobacterium*. Each column displays the E-test (minimum inhibitory concentration in µg/ml) and disc diffusion (inhibitory zone diameter in mm) results for a tested antimicrobial, classified based on the presence of target resistance determinants (*tetO, ermX*, and *gyrA* S115 and/or D119 mutation).

## Discussion

Our study is among the first to use the combination of anaerobic microbiology and genomics to study the diversity and antimicrobial susceptibility of a *Bifidobacterium* species in a developing country setting and identifies candidate therapeutic probiotics that may be relevant for Southeast Asia. We found that the *B. pseudocatenulatum* population within each individual is distinct and diverse. Intra-clonal variation in the pangenome usually stems from the gains or losses of glycosyl hydrolases and associated carbohydrate transporters, thus creating divergent metabolic functions, even for isolates within the same evolving clone. Our results reiterate previous findings on the genomic characterization of the *Bifidobacterium* genus (6) and *Bp* in the European population (33), showing that the species harbors an expansive repertoire of enzymes (GH13, GH43) responsible for the assimilation of complex plant-derived carbohydrates, but not host-derived glycan (mucin, HMO). In line with this observation, experimental study showed that *Bp* could utilize several components of arabinoxylan hydrolysates (AXH), via the upregulation of three GH43 – ABC transporter clusters (34). This feature likely explains the abundance and persistence of *Bp* through adulthood in the Vietnamese gut microbiota, since the organism may thrive on non-digestible carbohydrates enriched with fruits and vegetables.

We found that enzymes homologous to *B. longum* α-L-arabinofuranosidases (ArafA, ArafB, and ArafE) are potentially more abundant in adult-derived *Bp*, which reflects the adaptation of the species to the more complex fiber-rich diet usually present in adulthood. Similarly, a micro-evolution study of *B. longum subsp. longum* in Japan highlighted the enrichment of these homologs in strains derived from an elderly population (12). The above evidence could suggest a pattern of convergent evolution across different *Bifidobacterium* species, indicative of adaptation to resource availability. As these enzymes were predicted to be secreted in our *Bp* isolates, they may facilitate cross-feeding pathways with other bacteria incapable of utilizing complex arabinoglycan hydrolysates. Metabolic cross-feeding has been noted in *Bifidobacterium*, in which the fermentation end-products lactate and acetate can be utilized by anaerobes *Eubacterium hallii* (35) and *Faecalibacterium prausnitzii* (36), respectively, to produce butyrate. Likewise, *Bp* capable of degrading HMO were shown to release simpler metabolites, which supported the growth of other *Bifidobacterium* strains from the same breast-fed individual (33).

The EPS biosynthesis region was present in all recovered *Bp* genomes; this region is subjected to a high degree of genetic modification. The genetic composition of the EPS region is potentially more diverse in child-derived *Bp*, possibly owing to a greater extent of HGT in the *Bifidobacterium*-predominant gut microbiota in children. Previous studies have found that the EPS structure is critical for elucidating the *Bifidobacterium*-host interaction. In a simulated human intestinal environment, genes related to EPS biosynthesis were substantially up-regulated (37). The surface EPS grants protection against low pH and bile salt in the gastrointestinal environment (38, 39). Moreover, the presence of EPS in *B. breve* was associated with lower proinflammatory cytokines and antibody responses, which facilitate its persistent colonization in mouse models (38). However, it is known that *Bifidobacterium* with different EPS structures, even within a single species, can elicit differing immunological responses *in vitro* (40). For example, high-molecular-weight EPS is more likely to induce weaker immune responses, potentially because these encapsulating structures shield the complex protein antigens on the bacterial surface from interaction with immune cells (41). Furthermore, a specific EPS from *B. breve* has been shown to be metabolized by some members of the infant gut microbiota, indicating that EPS further facilitates cross-feedings between gut bacteria (42). Therefore, the considerable diversity shown by different EPS genomic characterizations in our *Bp* collection suggests the induction of varying and strain-dependent immunological responses within the human host. Recently, rhamnose-rich EPS were shown to elicit a moderate secretion of pro-inflammatory cytokines, prompting a mild boost to innate immunity (43). The EPS genomic clusters of the well-researched probiotic CECT7765, which has demonstrated a variety of health-promoting traits (19, 26), and several of our *Bp* isolates carried the *rmlABC* locus responsible for rhamnose biosynthesis. Future studies should investigate whether different rhamnose-rich EPS in *Bifidobacterium* confer similar impacts on inflammatory responses and ultimately on host health.

Our findings concur with previous studies on AMR in *Bifidobacterium*, showing that the carriage of *ermX* (44) and *tet* variants (45, 46) is common in the genus. These elements induce decreased susceptibility to azithromycin and tetracycline, respectively, which we confirmed in our study and has been observed in previous investigations (47). We also report a high prevalence of metronidazole resistance in these Vietnamese *Bp*. Metronidazole resistance in *Bifidobacterium* has been observed occasionally (48) and was recently reported in *Bp* causing pyogenic infections (49), but the resistance mechanism remains elusive. In addition, we showed here that mutation(s) in the QRDR of *gyrA* (S115 and/or D119) are likely to stimulate increased resistance to ciprofloxacin in *Bifidobacterium*. The encompassing region SAIYD (position 115 – 119 in GyrA) in wildtype *Bp* corresponds to the conserved SA[**X]**YD (83 – 87) in *Escherichia coli*’s GyrA, the most well-studied QRDR (32). However, unlike *E. coli*, which requires triple mutations (two in *gyrA*, one in *parC*) for full ciprofloxacin resistance (≥2 µg/mL), a single mutation in *Bp* may elevate ciprofloxacin MIC to ≥32 µg/mL (>16 fold increase). The ease of such a single-step process may explain the high degree of independent resistant mutations across different clones. Ciprofloxacin, azithromycin, and metronidazole are frequently prescribed for treatment of various infections in Vietnam. Prolonged exposure to these antimicrobials could induce resistance in *Bifidobacterium*, facilitated by the mobility of resistance determinants in the gut microbiota environment (46, 50). Though several studies have investigated the disturbance and recovery of the gut microbiome post-antibiotic treatment (51, 52), it is unknown how AMR in specific gut commensals (i.e. *Bifidobacterium*) impacts upon these ecological trajectories. Multi-drug resistance, which was observed in some recovered *Bp* may translate into higher survivability during a course of respective antimicrobial treatment, accelerating the recovery of the gut microbiota. Further studies are needed to test this hypothesis, especially given the potential high prevalence of AMR in *Bifidobacterium* in developing country settings.

There are limitations to our study. Since the study design was cross-sectional, we were not able to investigate the micro-evolution and population structure of *Bifidobacterium* within each participant over time. Our study is limited to genomic profiling, so further work is needed to validate *in-silico* predictions related to carbohydrate utilization and the EPS interaction with host cells. These drawbacks notwithstanding, we have a detailed collection of Vietnamese *Bp*, from which a probiotic candidate could be developed. This approach allowed us to tap into the diversity of resident *Bifidobacterium* within an indigenous population, for whom the probiotics or microbiome-targeted complementary foods will benefit. It also ensures that the candidates are well-tolerated and compatible with the local food matrix, maximizing its likelihood to colonize the target population. For instance, *Lactiplantibacillus plantarum* ATCC 202195, isolated from an Indian infant, was shown to colonize the neonatal gut for up to four months when orally administered with fructooligosaccharides (FOS) (53). This synbiotic (*L. plantarum* + FOS) proved successful in reducing the incidence of sepsis and death in rural Indian neonates, according to results published in a large scale randomized trial (54). Certain additional features may be considered when selecting a *Bp* candidate for development, including immunomodulatory potential (EPS similarity to other efficacious probiotics), metabolic flexibility (extensive glycosyl hydrolase content), survivability under antimicrobial pressures (multi-drug resistance), and contribution to overall gut health (production of acetate)(4). Thorough understanding on *Bp*’s metabolic capability allows for rational synbiotic design, which optimizes the bacterial survival and colonization in the gut. Regarding AMR, strains with resistance to metronidazole (epigenetic regulation) and/or ciprofloxacin (QRDR mutations) are more fitting as these resistance determinants are not transferrable to other members of the gut microbiota. Our study represents the primary step in identifying and characterizing potential bespoke probiotics from a developing country. This approach could be applied to develop microbiome-mediated therapeutics for other conditions in alternative locations, which will tap into the diversity and functionality held within the gut microbiota of those residing in LMICs, an area which is currently underexplored.

## Materials and Methods

### Study design and sample collection

Samples in this study originated from a cross-sectional study (from May to June 2017) to investigate the gut microbiomes of a healthy Vietnamese population. The study recruited healthy Vietnamese adults aged 18 to 60, who were at the time employed at the Oxford University Clinical Research Unit (OUCRU), Ho Chi Minh City, Vietnam and who were a parent or guardian of a child aged 9 to 60 months for whom they also consented to be enrolled in this study. Written informed consent was obtained from the adult participants and from the parent/guardian on behalf of child participants. Participants who had used antimicrobials in the three months prior to recruitment, or those who have or were recovering from chronic intestinal disease, chronic autoimmune disease or allergies were excluded from the study. Adults who experienced gastrointestinal infections in the last six months were also excluded. Based on these exclusion criteria, eligibility for study participation was self-assessed by the study participants. Recruitment was coordinated by a sample manager who ensured participant and specimen anonymity to other study staff. The study eventually recruited 21 adult and 21 child participants. Ethical approval for this study was obtained from the Oxford Tropical Research Ethics Committee (OxTREC, ID: 505-17).

Stool samples were collected from participants using non-invasive procedures. Briefly, participants were provided with a Protocult Collection Device (Ability Building Center, USA), including a transport container and a Ziploc bag. Specimens were labelled (with participant initials and date of collection) by the participants or their parent/guardian and stored in the freezer until delivery to OUCRU laboratories.

### Shotgun metagenomic sequencing and analysis

Total DNA extraction was performed on freshly collected stool samples (n=42) using the FastDNA Spin Kit for Soil (MP Biomedicals, USA) following the manufacturer’s procedures. These include a bead-beating step on a Precellys 24 homogenizer (Bertin Instruments, France). All DNA samples were then shipped to the Wellcome Trust Sanger Institute (WTSI, Hinxton, United Kingdom) for shotgun metagenomic sequencing on the Illumina HiSeq2000 platform. All output sequencing libraries were subjected to and passed the quality check on the WTSI pipeline. Taxonomic profiling was performed using the read-based Kraken approach (55) on a curated database of human gut microbial genomes, which include representative genomes (to species level) from the RefSeq database and ones sequenced from the collection in the Lawley lab (WTSI) (56).

### *Bifidobacterium* culture and identification

Samples with reads attributed to *Bifidobacterium pseudocatenulatum* (Bp) above 0.1% of total sequenced reads (7 adults, 9 children) were subjected to *Bifidobacterium* anaerobic culturing, using a Whitley A35 anaerobic workstation (Don Whitley Scientific, United Kingdom) containing 5% CO_2_, 10% hydrogen and 85% nitrogen gas. Briefly, fecal samples were homogenized in PBS (0.1 g stool per ml of PBS), and several ten-fold dilutions (10^−4^ to 10^−7^) were plated onto de Man Rogosa and Sharpe (MRS) media (BD Difco, USA) supplemented with 50 mg/L of mupirocin (PanReac AppliChem, Germany) and L-cysteine.HCl (Sigma-Aldrich, Germany) (13). Plates were incubated at 37^0^C for 48 hours, and ∼10 different colonies were randomly picked from each fecal sample and re-streaked on new MRS plates to confirm purity. For each of these individual bacterial isolates, taxonomic identities were confirmed on the matrix assisted laser desorption/ionization time-of-flight mass spectrometer (MALDI-TOF, Bruker). In addition, each isolate was subjected to full-length 16S rRNA gene PCR and capillary sequencing, using the primer pair 7F (5’-AGAGTTTGATYMTGGCTCAG-3’) and 1510R (5’-ACGGYTACCTTGTTACGACTT-3’) (56). *Bifidobacterium* species identity was confirmed by comparing the output sequence with the NCBI 16S rRNA gene database. In total, 185 isolates were confirmed as *Bifidobacterium* species, of which 49 were *B. pseudocatenulatum* (as identified by both methods).

### Whole genome sequencing, pangenome analysis and phylogenetic inference of *Bifidobacterium pseudocatenulatum*

Forty-nine confirmed *Bp* isolates were subjected to DNA extraction using the Wizard Genomic Extraction Kit (Promega, USA). For whole genome sequencing (WGS), one nanogram of extracted DNA from each isolate was input into the Nextera XT library preparation kit to create the sequencing library, as per the manufacturer’s instruction. The normalized libraries were pooled and then sequenced on an Ilumina MiSeq platform to generate 250bp paired-end reads.

The sequencing quality of each read pair was checked using FASTQC (57), and Trimmomatic v0.36 was used to remove sequencing adapters and low quality reads (paired end option, SLIDINGWINDOW:10:20, MINLEN:50) (58). For each trimmed read set, *de novo* genome assembly was constructed separately using SPAdes v3.12.0, with the error correction option and default parameters (59). Annotation for each assembly was performed using Prokka v1.13, using input of other well-annotated *Bifidobacterium* sequences as references (60). The pangenome of 49 sequenced *B. pseudocatenulatum*, together with public references of the species (DSM20438, CECT_7765, and five assembled genomes from a micro-evolution study in China (15)), was determined using panX with default settings (61). In brief, panX reconstructs individual gene trees, and uses these in an adaptive post-processing step to scale the thresholds relative to the core genome diversity, instead of relying on a specific single nucleotide identity cut-off. The resulting core-genome (1116 single-copy genes, 137,285 SNPs) was input into RAxML v8.2.4 to construct a maximum likelihood phylogeny of all 56 queried genomes, under the GTRGAMMA model with 500 rapid bootstrap replicates (62). This phylogeny delineates the separation of two lineages, major (n=52) and minor (n=4).

To accurately identify the taxonomic identity of the four isolates belonging to the minor lineage (C01_D5, C01_H5, C01_C5, and C02_A8), we used panX and RAxML as described above to infer the core-genome phylogeny of these four isolates together with several *Bifidobacterium* references. These include *B. adolescentis* 15703, *B. pseudocatenulatum* (DSM20438, 12), *B. catenulatum* (MC1, BCJG468, 1899B, DSM16992, HGUT, BIOMLA1, BIOMLA2, JG), and *B. kashiwanohense* (JCM15439, APCKJ1, PV20-2). For the remaining 52 strains, we mapped each read set to the reference DSM20438, using BWA-MEM with default parameters, and SNPs were detected and filtered using SAMtools v1.3 and bcftools v1.2 (63). PICARD was used to remove duplicate reads, and GATK was employed for indel realignment, as previously recommended (64). Low quality SNPs were removed if they matched any of these criteria: consensus quality < 50, mapping quality < 30, read depth < 4, and ratio of SNPs to reads at a position < 90%. Bedtools v2.24.0 was used to summarize the mapping coverage at each position in the reference (65). A pseudo-sequence (with length equal to that of the mapping reference) was created for each isolate to integrate the filtered SNPs, region of low mapping coverage, and invariant sites, using the vcf2fa python script (--min_cov=4, https://github.com/brevans/vcf2fa). Together with the mapping reference, pseudo-sequences were concatenated to create an alignment, which was then input into ClonalFrameML (branch extension model; kappa = 9.305, emsim=100, embranch dispersion = 0.1) to remove regions affected by recombination (66). This created a SNP alignment of 10,716 bp, which served as input for RAxML to infer a maximum likelihood phylogeny of 45 *sensu stricto Bp*, under the GTRGAMMA model with 500 rapid bootstrap replicates (5 iterations). In addition, seven isolates belonging to the C04_cluster were subjected to mapping (to DSM20438) and SNP calling, using the aforementioned parameters. The resulting alignment was input into Gubbins to remove regions of recombination (67), followed by maximum likelihood reconstruction using RAxML.

### Determination of carbohydrate-active enzymes and antimicrobial resistance determinants

Representatives from each gene family (n=4,333), as identified in the pangenome by panX as described above, were input into the dbCAN2 metaserver (http://bcb.unl.edu/dbCAN2/blast.php) to annotate genes involved in carbohydrate utilization (68). These carbohydrate active enzymes (CAZymes) include glycosyl hydrolases (GH), glycosyl transferase (GT), glycosyl lyase and esterase. A candidate gene was considered a CAZyme if the dbCAN2 output returned any positive hits from the three detection algorithms: HHMER, Hotpep, and DIAMOND. In addition, assembled genomes were screened by ARIBA against a curated resistance determinant database (ResFinder) to detect the presence of acquired resistance genes (--nucmer_min_id = 95, --nucmer_min_len = 50) (69, 70). QRDR mutations were manually screened by aligning the *gyrA, gyrB, parC*, and *parE* homologs of sequenced genomes, retrieved from the constructed pangenome analysis.

### Antimicrobial susceptibility testing of *Bifidobacterium*

For each PC, we selected one representative isolate for antimicrobial susceptibility testing, except for the case of C16_cluster_1, in which two isolates were included because they showed different genetic composition in the EPS biosynthesis cluster (17 *B. pseudocatenulatum* and 2 *B. catenulatum*). Four control strains were also included: *B. pseudocatenulatum* DSM20438, *B. longum subsp. longum* NCIMB 8809, *Staphylococcus aureus* ATCC 25923, and *S. aureus* ATCC 29213. Strains were maintained in Brain Heart Infusion (BHI) with 20% glycerol at −80^0^C prior to resuscitation on MRS and Luria-Bertani agar (Oxoid, UK), for *Bifidobacterium* and *S. aureus*, respectively. LSM-cysteine formulation (90% Iso-Sensitest broth [Oxoid, UK] and 10% MRS broth, supplemented with 0.3g/l L-cysteine.HCl, with pH adjusted to 6.85 ± 0.1) was chosen for antimicrobial susceptibility testing of *Bifidobacterium*, as recommended previously (71). Muller-Hinton (MH) media was chosen for testing of *S. aureus*. Prior to testing, strains were pre-cultured on LSM-cysteine agar (48 hours for *Bifidobacterium*) or MH agar (24 hours for *S. aureus*) under the specified incubation conditions.

For the disc diffusion method, inocula were prepared by suspending *Bifidobacterium* colonies from LSM-cysteine plates into 5ml of 0.85% NaCl solution (adjusted to McFarland standard 1), which were then spread onto LSM-cysteine plates (72). Subsequently, antimicrobial discs (Biomeriux, France), including ceftriaxone (30µg), ciprofloxacin (5µg), tetracycline (30µg), amoxicillin/clavulanic acid (30µg), azithromycin (15µg), and metronidazole (5µg) were applied. Plates were incubated under anaerobic conditions for 48h at 37^0^C, followed by measurement of the diameters of the inhibition zones, including the diameter of the disc (mm). For each isolate, the procedures were repeated five times to evaluate day-to-day reproducibility of the method, with reproducibility defined as percentage of samples within ±4 mm variation in zone diameter (72). For E-tests, inocula were prepared as described above, and the resuspended solution was spread onto LSM-cysteine plates. Plates were left to dry for ∼15 minutes, after which E-test strips were applied (ceftriaxone, ciprofloxacin, tetracycline, amoxicillin/clavulanic acid, azithromycin, and metronidazole; Biomeriux, France). The MIC (µg/ml) was assessed after 48-hours incubation, with MIC defined as the value corresponding to the first point on the E-test strip where growth did not occur along the inhibition ellipse.

### Data analysis and visualization

All data analyses were conducted in R (73) using multiple packages, including ggplot2 and ggtree for visualization (74, 75). To compare the accessory gene content between child- and adult-derived *Bp*, we selected a representative genome from each of the identified phylogenetic clusters (PCs) in the *Bp* phylogeny (Figure 2, n=16). We also included one additional genome for PCs showing high intra-clonal variation in the accessory genome (A05_cluster_1, A10_cluster, C04_cluster, C16_cluster_1, C16_cluster_2). This resulted in a set of 21 independent genomes (adult: 10, child: 11). Only gene families (as identified by panX) that are present in five to sixteen genomes (n=606) were considered for statistical testing (Fisher’s exact test) to investigate the differences in genetic composition of the two groups (child vs. adult). Due to the limited number of tested genomes, correction for multiple hypothesis testing was not employed, and we reported candidates with p value ≤ 0.05 as indicating potential differences.

Artemis and Artemis Comparison Tool (ACT) were utilized to visualize the presence of selected genetic elements in the genomes (76). The EPS biosynthesis cluster was defined as a genomic region flanked by the priming glycosyltransferase *rfbP* or *cpsD*, and encompassing several GTs, polysaccharide export *rfbX*, oligosaccharide repeat unit polymerase, tyrosine kinase, and tyrosine phosphatase (37, 41). This region was extracted from targeted *Bifidobacterium* genomes, and queried against the NCBI public database using BLASTN to identify the most similar variants. Comparisons between different EPS regions were visualized by Easyfig (77).

## Acknowledgements

The authors wish to thank all participants and their parents/guardians for their participation in the study, and Dr. Magdalena Kujawska for her advice in *Bifidobacterium* culturing.

## Data availability

Raw sequence data are available in the NCBI Sequence Read Archive (project PRJNA720750: Genomic diversity of *Bifidobacterium pseudocatenulatum* in the Vietnamese population).

